# Emergency hematopoiesis proceeds without contribution of hematopoietic stem cells

**DOI:** 10.1101/2022.10.05.510973

**Authors:** Clara M. Munz, Nicole Dressel, Minyi Chen, Tatyana Grinenko, Axel Roers, Alexander Gerbaulet

## Abstract

Hematopoietic stem cells (HSCs) are the ultimate source of blood and immune cells. Under homeostatic conditions, these cells are considered a quiescent reserve population. However, it is not clear to what extent HSCs participate in emergency responses. Herein, we use fate mapping and proliferation tracking mouse models, which cumulatively record HSC activity *in situ*. We observed no direct contribution of HSCs to mature blood cell regeneration in response to common hematopoietic emergencies, including inflammation or blood loss. Innate immune training, in which HSCs were proposed to store and integrate information on previous infections, did not alter HSC activity upon secondary exposure. Only severe myeloablation resulted in a robust increase of HSC contribution. Our data demonstrates that HSCs do not directly participate in the regeneration of mature blood cells and therefore do not represent a reserve population to compensate for physiological hematopoietic perturbations.

## Introduction

The hematopoietic system continuously generates enormous numbers of mature blood cells (Cosgrove et al., 2021) and its output can be substantially accelerated by blood loss or infections. Hematopoietic stem cells (HSCs), which reside atop the hematopoietic hierarchy, give rise to mature blood cells via diverse progenitor intermediates. Upon transplantation, only this somatic stem cell population is able to reconstitute all blood cell lineages lifelong, which is the basis for therapeutic or experimental HSC transplantation (Purton and Scadden, 2007; Thomas et al., 1957). While post-transplantation hematopoiesis is driven by few HSC clones (Naik et al., 2013; Rodriguez-Fraticelli et al., 2020; Sun et al., 2014), fate tracking of native hematopoiesis in adult mice revealed only rare, but continuous and polyclonal HSC differentiation (Bowling et al., 2020; Busch et al., 2015; Pei et al., 2017; Sun et al., 2014). Moreover, murine HSCs remain largely quiescent in adult steady-state hematopoiesis (Visser et al., 1981; Wilson et al., 2008). A recent study revealed the enormous regenerative capacity of progenitor populations, which despite lack of long-term transplantation potential, contribute lifelong to native hematopoiesis (Patel et al., 2022). Models of HSC depletion further emphasize the robustness of blood formation in the absence of HSCs (Schoedel et al., 2016; Sheikh et al., 2016). Currently, HSCs are perceived as a reserve population, which is essential to boost blood cell output in situations of acute demand, as for example in response to infection or blood loss (King and Goodell, 2011; Takizawa et al., 2012; Trumpp et al., 2010). HSCs are activated via pattern recognition and cytokine receptors and participate in hematopoietic stress responses, ultimately resulting in a gradual loss of their transplantation potential (reviewed by Caiado et al., 2021). Importantly, innate immune training of hematopoiesis (Netea et al., 2020), i.e. a memory of the hematopoietic system allowing for faster and more efficient innate immune responses to a recurring infectious stimulus, was reported to be epigenetically imprinted in HSCs (de Laval et al., 2020; Li et al., 2022).

Here, we used mouse models of fate and proliferation tracking to study HSC activity during hematopoietic stress. We found that innate immune signaling resulting in massively enhanced blood cell output was reflected in little, if any, increase in HSC contribution. Similarly, hematopoietic stress caused by the depletion of mature blood cells did not trigger accelerated stem cell proliferation or differentiation. Only pharmacological or irradiation-induced severe ablation of hematopoietic progenitors triggered a robust plus in HSC activity. We thus argue against HSCs constituting a reserve population that compensates hematopoietic emergencies.

## Results

### Fate mapping and proliferation tracking of perturbed hematopoiesis

To investigate the contribution of HSCs to hematopoietic stress responses, we employed the *Fgd5*^ZsGreen:CreERT2^/*R26*^LSL-tdRFP^ fate mapping mouse model, in which tamoxifen (TAM) selectively induces inheritable RFP expression in adult immuno-phenotypic (lin^-^ Sca-1^+^ Kit^+^ (LSK) CD48^-/lo^ CD150^+^) HSCs (Figures 1A-C and S1) (Morcos et al., 2022). This model preferentially labels a subpopulation of HSCs at the apex of the hematopoietic hierarchy with high transplantation potential (Gazit et al., 2014), identified by high surface expression of CD201 (EPCR) and Sca-1 (termed “ES HSCs” or “tip” HSCs). To monitor the propagation of label from tip HSCs to their progeny and to adjust for the inter-individual variability of labeling efficiency, we calculated the fraction of ES HSC-derived label in each cell population under investigation. This approach revealed that ES HSCs constantly gave rise to HSCs, evidenced by equilibration of their label within the first year of life (Figure 1D) and revealing that the system initially indeed labeled ‘tip’ HSCs (Morcos et al., 2022; Takahashi et al., 2021). In addition to HSC label propagation, we determined the divisional history of hematopoietic stem and progenitor cells (HSPCs) employing *R26*^rtTA^/*Col1A1*^H2B-GFP^ mice (Figure 1E) (Foudi et al., 2008; Kanda et al., 1998). Doxycyclin (DOX) administration to these mice induces ubiquitous H2B-GFP labeling. After withdrawal of the inducer, each cell division halves H2B-GFP intensity until background fluorescence levels are reached (Morcos et al., 2020) (Figure 1F).

**Figure 1:**
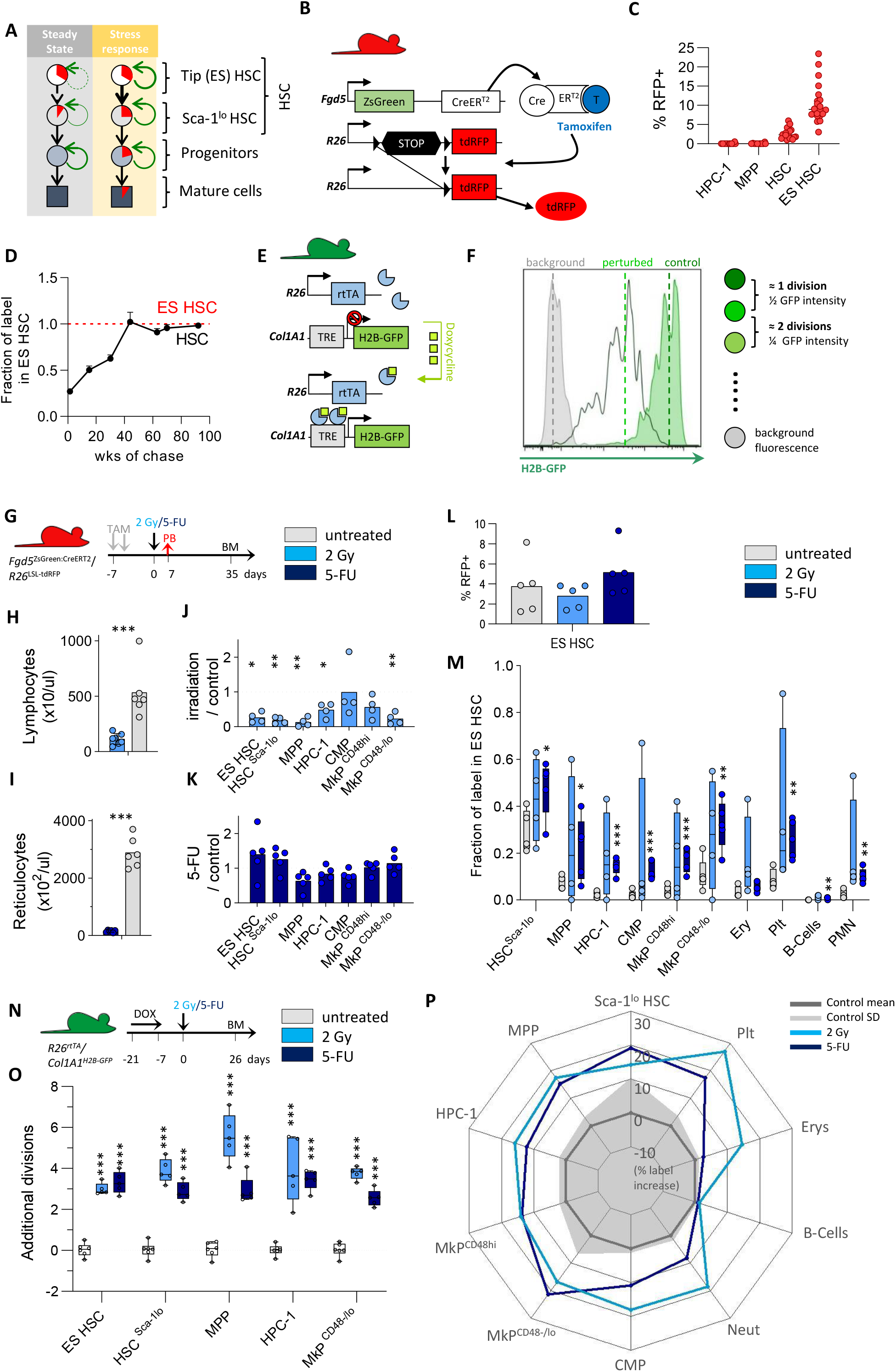
Fate mapping and proliferation tracking of perturbed hematopoiesis. **(A)** How perturbations of hematopoiesis alter differentiation (red) and proliferation (green) of HSPCs will be studied. **(B)** *Fgd5*^ZsGreen:CreERT2^/*R26*^LSL-tdRFP^ mouse model for fate mapping of HSCs. **(C and D)** RFP labeling of HSPCs isolated from TAM-induced *Fgd5*^ZsGreen:CreERT2^/*R26*^LSL-tdRFP^ mice. Data from Morcos et al. (2022) (**C**) Initial labeling frequencies of BM HSPCs 10 days after induction. **(D)** Labeling of total HSCs relative to ES HSCs (dotted line, 8-20 mice/timepoint). **(E)** *R26*^rtTA^/*Col1A1*^H2B-GFP^ mouse model for proliferation tracking of HSPCs. (**F)** Representative H2B-GFP histograms of ES HSCs isolated from 5-FU-perturbed and control mice (see also Figure S2C-E). **(G-M)** *Fgd5*^ZsGreen:CreERT2^/*R26*^LSL-tdRFP^ mice were TAM-induced and perturbed with either 2 Gy γ-radiation (n=4), 5-FU (n=5) or left untreated (n=5). (**G**) Experiment scheme; **(H)** PB lymphocyte count 7 d after irradiation; **(I)** PB reticulocyte numbers 7 d after 5-FU; **(J-K)** Ratios of relative BM compartment sizes (% cells among total lin^-^ BM cells) between perturbed and control (dotted line) animals. **(L)** Percentages of RFP^+^ ES HSCs. **(M)** Fraction of RFP-labeled cells relative to ES HSCs. **(N and O)** *R26*^rtTA^/*Col1A1*^H2B-GFP^ animals were DOX-pulsed and exposed to either 2 Gy (n=5), 5-FU (n=5) or left untreated (n=6). (**N**) Experiment scheme; (**O**) numbers of additional divisions in response to perturbation (see also Figures S2C-E). **(P)** Transformation of data shown in **M** to visualize the net effects of myeloablation on label propagation. Mean (dark grey line) and variance (SD, shaded area) of control mice (n=41 from 8 independent experiments) are shown (see also Figures S2F-H).

Immuno-phenotypes of HSPCs activated by inflammatory stimuli are poorly defined, as signaling under stress conditions perturbs surface marker expression (Bujanover et al., 2018; Kanayama et al., 2020) (Figure S2A-B). However, both reporter mouse models record either cell division (H2B-GFP dilution) or HSC contribution (RFP label propagation) in a cumulative fashion and can be read out after the acutely perturbed hematopoietic system has returned to steady state and faithful marker expression is restored.

### Myeloablation stimulates HSC activity

The hematopoietic system is particularly susceptible to ionizing radiation or chemotherapy, which rapidly deplete cycling HSPCs and are commonly referred to as myeloablation. To investigate the contribution and proliferation of HSCs in response to this perturbation, we subjected previously induced *Fgd5*^ZsGreen:CreERT2^/*R26*^LSL-tdRFP^ as well as *R26*^rtTA^/*Col1A1*^H2B-GFP^ reporter mice to either 2 Gy γ-radiation or 5-fluorouracil (5-FU) (Figures 1G-P and S2C-H). To control for successful myeloablation, peripheral blood (PB) analysis 7 days later revealed a profound reduction of lymphocytes or reticulocytes, respectively (Figures 1H, I). We analyzed the composition of the lineage-negative (lin^-^) bone marrow (BM) compartment 5 weeks after perturbation and found a significant reduction of HSCs and MPPs after irradiation, while all compartments of 5-FU exposed BM appeared normal (Figures 1J, K). ES HSCs represent the ultimate source of propagated label and therefore serve as the reference for normalization. Myeloablation did not alter the RFP labeling of ES HSCs (Figure 1L), but strongly accelerated label propagation from ES HSCs to more differentiated Sca-1^lo^ HSCs (Morcos et al., 2022; Morcos et al., 2017), progenitors and, to a lesser extent, to mature blood cells (Figure 1M). To extract the net effect of perturbation on proliferation, we calculated additional divisions of HSPCs in myeloablated animals compared to controls. This revealed about 3 additional divisions in HSCs including ES HSCs in response to both perturbations (Figures 1N, O and S2C-E). γ-radiation resulted in massive proliferation of MPPs and HPC-1s, thereby diluting the H2B-GFP label to the range of background controls and likely exceeding the maximum number of ^~^4-6 traceable divisions. Likewise, we calculated the net effect caused by perturbation on label propagation by subtracting the labeling of controls from the labeling of perturbed animals and plotted this data as a radar chart (Figures 1P and S2F-H). Taken together, our models faithfully reported the accelerated proliferation and differentiation of HSCs expected upon myeloablation (Bowling et al., 2020; Busch et al., 2015; Wilson et al., 2008).

### Rare contribution of tip HSCs to emergency myelopoiesis

While myeloablation proved a potent, but rather artificial stimulus for HSC activation, innate cytokine signaling during, for example, infection, represents a more physiological hematopoietic stress response. Lipopolysaccharide (LPS) is a major molecular pattern of gram-negative bacteria and triggers the expression of inflammatory cytokines via activation of Toll-like receptor (TLR) 4 (Beutler, 2009). HSCs were reported to directly sense LPS, thereby entering the cell cycle to initiate emergency myelopoiesis and LPS exposure *in vivo* caused loss of HSC transplantation potential (reviewed by Caiado et al., 2021). We intraperitoneally injected TAM-induced *Fgd5*^ZsGreen:CreERT2^/*R26*^LSL-tdRFP^ mice with a single dose of LPS, and demonstrated induction of emergency myelopoiesis by recruitment of neutrophils to the injection site and increased interleukin (IL)-6 levels in PB plasma (Figures 2A-C). Unexpectedly, the massive acceleration of blood cell output upon LPS exposure was not reflected in faster label propagation from ES-HSCs throughout the hematopoietic system (Figure 2D, S3A-C). While there was a slight label increase in Sca-1^lo^ HSCs, MPPs and common myeloid progenitors (CMPs), labeling of mature erythrocytes and leukocytes did not increase, demonstrating that label from tip HSCs does not reach mature myeloid cell populations even within 2 weeks after LPS-induced emergency myelopoiesis. Consistent with HSC fate mapping, LPS injection did not induce robust proliferation of early HSPCs (Figure 2E, S3D).

**Figure 2:**
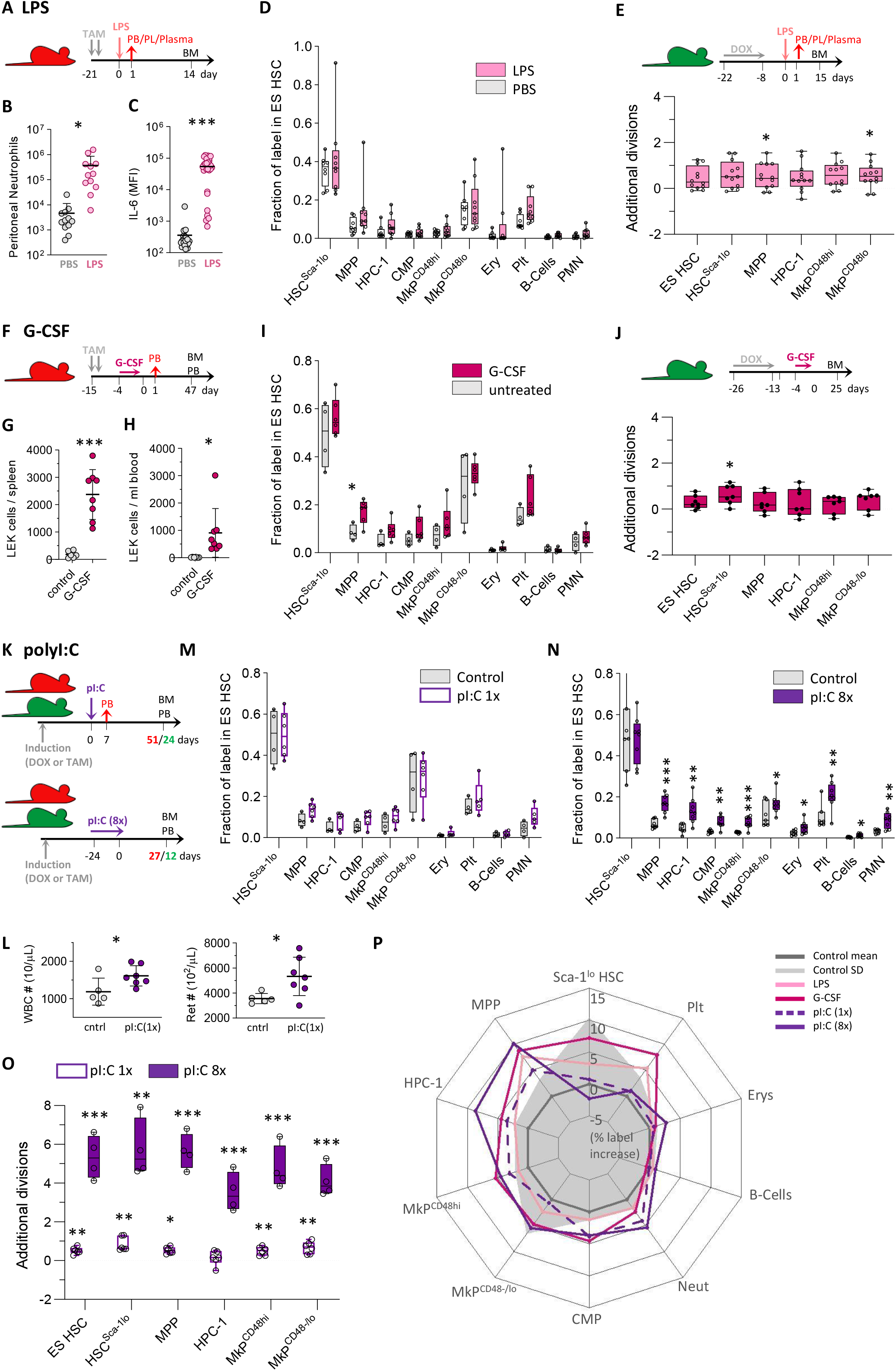
Limited contribution of HSCs to emergency myelopoiesis and type I IFN stimulation. **(A-D)** *Fgd5*^ZsGreen:CreERT2^/*R26*^LSL-tdRFP^ mice were TAM-induced and injected with LPS (n=8) or PBS (n=12). **(B)** Neutrophil count in the peritoneal cavity 24 h after LPS. **(C)** Plasma levels of IL-6 90 min after LPS. **(D)** Fraction of RFP-labeled cells (relative to ES HSCs) between LPS and saline-treated animals. Data from two independent experiments. **(E)** Labeled *R26*^rtTA^/*Col1A1*^H2B-GFP^ animals were i.p. injected with LPS (n=14) or PBS (n=13) and the average number of additional divisions in response to LPS was determined. Data from two independent experiments. **(F-J)** *Fgd5*^ZsGreen:CreERT2^/*R26*^LSL-tdRFP^ (**F, I,** n=4-6/condition) and *R26*^rtTA^/*Col1A1*^H2B-GFP^ (**G, H, J,** n=4-7/condition) mice were s.c. injected with G-CSF or PBS for five consecutive days. (**G-H)** Numbers of LEK HSPCs in spleen (**G**) or PB (**H**) of *R26*^rtTA^/*Col1A1*^H2B-GFP^ animals on d6. **(K-O)** *Fgd5*^ZsGreen:CreERT2^/*R26*^LSL-tdRFP^ (**K-N,** n=4-8/condition) and and *R26*^rtTA^/*Col1A1*^H2B-GFP^ (**O,** n=4-7/condition) mice were i.p. injected with pI:C or PBS following either a single (1x) or repetitive (8x) administration protocol. (**L**) PB leukocyte and reticulocyte numbers 7 d after 1x pI:C. **(P)** Net effects of LPS, G-CSF or pI:C perturbation on RFP label propagation (% label increase relative to ES HSCs, transformation and display of data as in Figure 1P).

Granulocyte-colony stimulating factor (G-CSF) induces myeloid differentiation, as well as HSPC mobilization (Greenbaum and Link, 2011) and G-CSF secretion by endothelial cells in response to systemic LPS administration was identified as a key event of emergency myelopoiesis (Boettcher et al., 2014). We injected previously labeled *Fgd5*^ZsGreen:CreERT2^/*R26*^LSL-tdRFP^ and *R26*^rtTA^/*Col1A1*^H2B-GFP^ mice for five consecutive days with G-CSF (Figures 2F-J and S3E-L) and demonstrated mobilization of BM HSPCs to spleen and PB (Figures 2G, H) on day 6. BM analysis 47 and 25 days after G-CSF administration, respectively, uncovered a pattern of RFP label propagation (Figures 2I and S3E-G) and HSPC proliferation (Figures 2J and S3H) which resembled our LPS-treated animals. Specifically, we found moderately increased labeling of MPPs, but HSC-derived label did not reach mature blood lineages. Again, G-CSF did not robustly stimulate additional divisions in HSCs and early progenitor cells (Figures 2J and S3H). Since HSPC egress from the BM to the periphery is a major effect of G-CSF (Greenbaum and Link, 2011) and has previously been linked to cell proliferation (Bernitz et al., 2017; Morrison et al., 1997), we investigated the divisional activity of HSPCs directly after mobilization (Figure S3I-L) and detected a similar divisional history (less than or equal to one additional division) of both mobilized and control HSCs analyzed in BM and spleen. In contrast, the comparison of PB and BM in mobilized animals revealed extensive proliferation of PB HSCs (Figure S3I-J), which expressed surface markers linked to proliferation and differentiation (Figure S3K, L), suggesting selective mobilization of less primitive HSCs into the blood stream.

### Type I IFN stimulates HSC divisions but little differentiation

Acute type I interferon (IFN) exposure stimulates cell cycle entry of quiescent HSCs. The synthetic double stranded RNA analogue polyinosinic:polycytidylic acid (pI:C) induces type I IFN and inflammatory cytokines via TLR3 activation and recapitulates key features of a viral infection. Repetitive pI:C application was shown to attrite HSC transplantation potential (reviewed by Demerdash et al., 2021). To induce acute or prolonged type I IFN signaling, we injected labeled fate mapping and proliferation tracking mice once or repetitively (8x) with pI:C (Figures 2K-O and S3M-T), which resulted in an increase of PB leukocytes and reticulocytes persistent until 7 days after injection (Figure 2L). HSC label propagation was not accelerated upon a single administration of pI:C (Figures 2M and S3O). Repetitive pI:C injections resulted in moderate, albeit significant increases of labeled cells in all analyzed progenitor and mature blood cell populations with the exception of more committed Sca-1^lo^ HSCs (Figure 2N and S3S). Importantly, pI:C robustly stimulated the mitotic activity of HSPCs in a dose-dependent manner, with at least 5 additional divisions on average in HSCs and MPPs upon repetitive pI:C application (Figures 2O and S3P, T). Taken together, acute inflammatory cytokine signaling mimicking bacterial or viral infections did not induce a robust contribution of HSCs to mature blood cells (Figure 2P). Only in response to prolonged type I IFN signaling, HSPCs vigorously proliferated, but compared to myeloablation, this resulted in much weaker propagation of tip HSC-derived label.

### Massive loss of RBC and Plt does not stimulate HSC output

Red blood cells (RBCs) constitute 90% of all blood cells and account for 80-90% of all daily renewed cells in mice and men (Sender and Milo, 2021). To model acute blood loss, we injected our labeled reporter mouse models with two doses of phenylhydrazine (PHZ, Figures 3A-E), which destroys RBCs by forming methemoglobin and resulted in halved hematocrit levels one day after PHZ application (Figure 3B). Despite loss of approximately 50% of RBCs, which equals ^~^7.5×10^9^ cells / mouse, complete regeneration was achieved by day 14 (Figure 3C). However, in PHZ-perturbed mice, we did not observe increased label propagation from ES HSC (Figure 2D, S4A-C) and HSPC proliferation was not accelerated (Figure 3E, S4D), which reveals that billions of additional RBCs were regenerated without significant input from HSCs.

**Figure 3:**
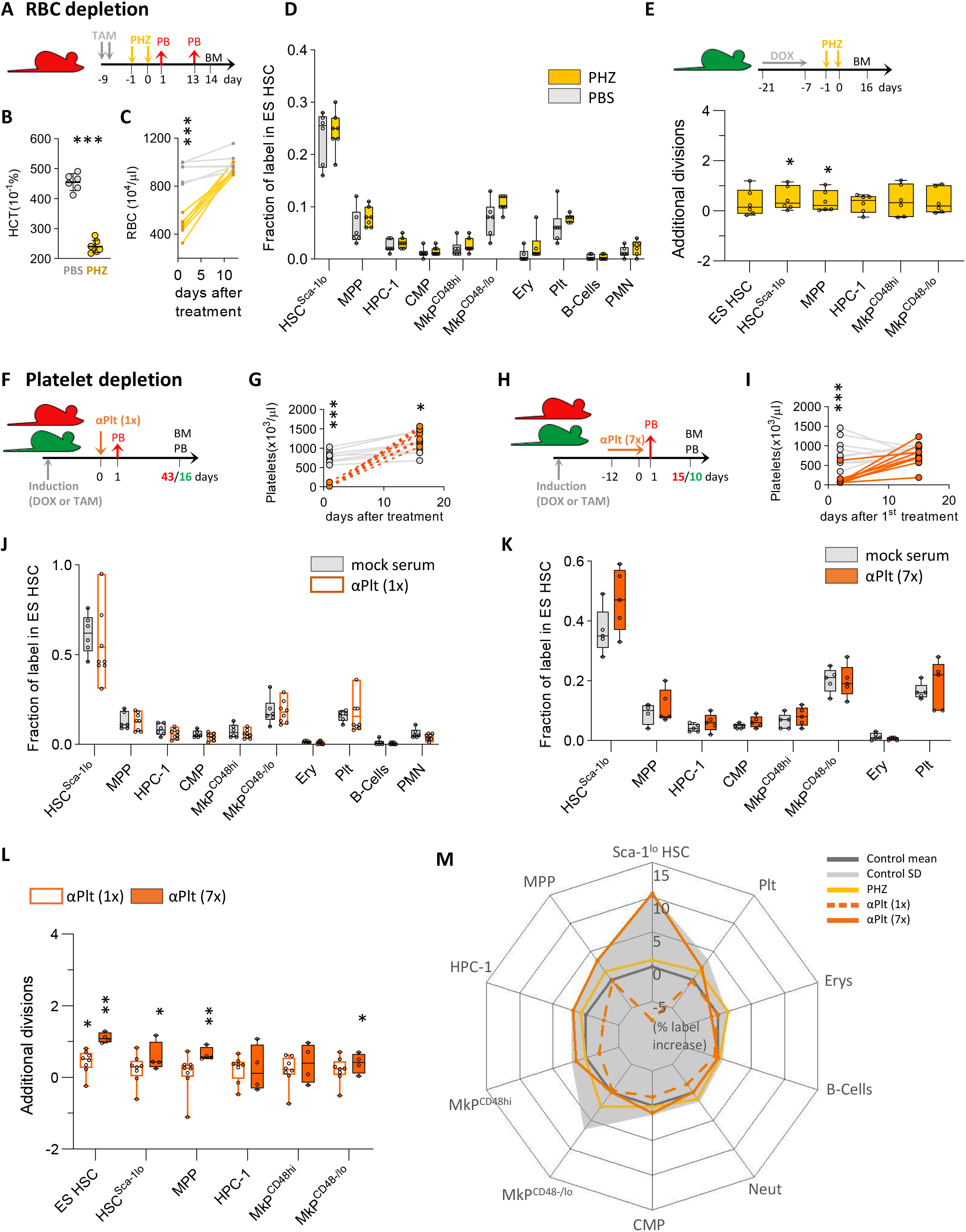
Limited contribution of HSCs to recovery from blood loss. **(A-E)** *Fgd5*^ZsGreen:CreERT2^*/R26*^LSL-tdRFP^ (**A-D,** n=6-7/condition) and *R26*^rtTA^/*Col1A1*^H2B-GFP^ (**E,** n=6-7/condition) mice were i.p. injected with phenylhydrazine (PHZ) or PBS. **(B)** Hematocrit (HCT) 24h after PHZ application and **(C)** PB RBC numbers. **(F-L)** *Fgd5*^ZsGreen:CreERT2^*/R26*^LSL-tdRFP^ (**F-K,** n=4-8/condition) and *R26*^rtTA^/*Col1A1*^H2B-GFP^ (**L,** n=4-8/condition) mice were treated with anti-platelet (αPlt) serum or control serum following a single (1x, **F, G, J, L**) or repetitive (7x, **H, I, K, L**) application protocol. (**G, I**) Platelet numbers in PB. **(M)** Net effects of RBC or PLT depletion on RFP label propagation (% label increase relative to ES HSCs; transformation and display of data as in Figure 1P).

Similar to RBCs, low platelet counts have potentially lethal consequences. To model acute as well as chronic thrombocytopenia, we depleted platelets (Plts) by either a single or repeated injection of anti-platelet (αPlt) serum (Figures 3F-I and S4E-L). Platelet numbers were completely recovered within 14 days after treatment (Figures 3G, I), but HSC-derived label propagation was not accelerated neither by acute nor by chronic platelet depletion (Figures 3J, K and S4G, K). αPlt serum dose-dependently induced less than or equal to a single additional division on average in early HSPCs (Figures 3L and S4H, L) as previously reported (Ramasz et al., 2019). Taken together, depletion of mature Plt and RBCs did not elicit substantial contribution of tip HSCs to mature blood cells (Figure 3M).

### Innate immune memory does not alter HSC proliferation and differentiation kinetics

Primary pathogen encounters were recently shown to induce long-lasting epigenetic adaptions of HSC to accelerate production of innate effector cells upon secondary exposure (Netea et al., 2020). Previous stimulation with β-glucan was reported to result in more rapid myeloid progenitor output upon subsequent exposure to LPS (Mitroulis et al., 2018). To investigate how innate immune training alters HSPC proliferation and differentiation, we exposed cohorts of our cell fate and proliferation reporter mouse models to the fungal cell wall component β-Glucan, followed by reporter induction and LPS stimulation 4 weeks later (Figures 4A-C, “trained”). Matched control groups did not receive the initial β-glucan training stimulus, but were exposed to LPS (Figures 4A-C, “untrained”). β-glucan administration caused a mild, but significant neutrophilia in PB 1 day later (Figure 4D). LPS administration resulted in a significant decline of PB Plt, eosinophils, neutrophils and lymphocytes (Figure 4E) within 24h, as well as in massive recruitment of neutrophils to the peritoneal cavity (Figure 4F), but all these effects of LPS were undistinguishable between trained and untrained mice. In addition, we determined plasma cytokine levels 90 min after LPS administration, however, β-glucan training resulted in non-significant alterations, trending towards higher IL-1β, −6, −10, tumor necrosis factor and MCP-1 production (Figures 4G and S4M). In both the fate and the proliferation tracking model, LPS stimulated some activity, however, this pattern was highly similar between trained and untrained animals (Figure 4H, I and S4N-Q). In summary, we show that β-glucan priming did not result in increased differentiation or proliferation of HSCs in response to subsequent LPS exposure.

**Figure 4:**
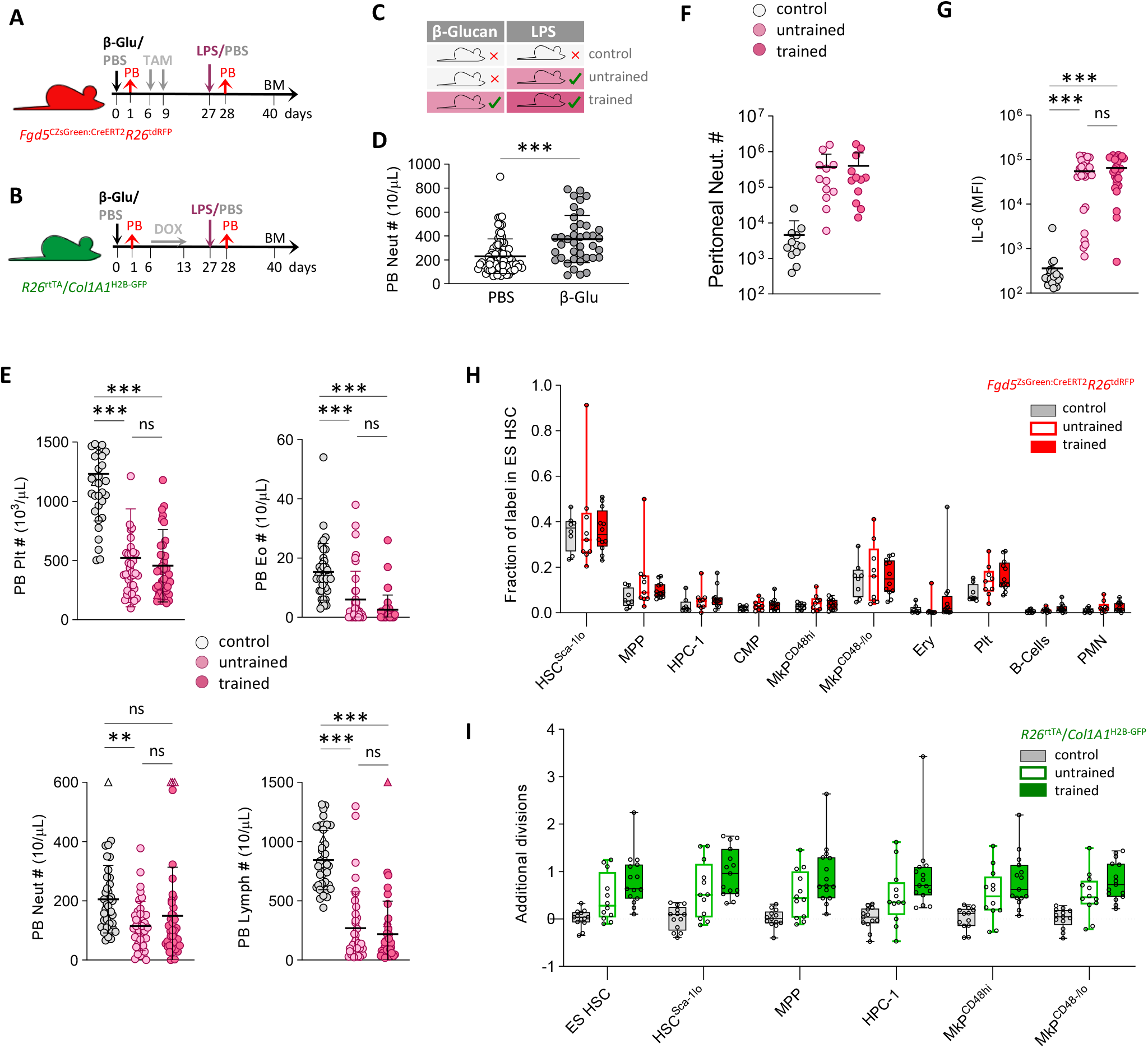
Innate immune memory does not alter HSC proliferation and differentiation kinetics. **(A-I)** *Fgd5*^ZsGreen:CreERT2^/*R26*^LSL-tdRFP^ (**A,** n=8-12) and *R26*^rtTA^/*Col1A1*^H2B-GFP^ (**B,** n=12-14) mice were either treated with PBS or β-glucan, followed by reporter induction, secondary treatment with LPS or PBS (experimental groups in **C**) and final PB and BM analysis. Data from two independent experiments. **(D-E)** PB neutrophil, lymphocyte, platelet or eosinophil numbers 24h after injection of β-glucan (**D**) or LPS (**E**). Data from 4 independent experiments. **(F)** Neutrophil numbers in the peritoneal cavity 24 h after LPS or PBS injection. **(G)** Levels of IL-6 in plasma isolated from control, untrained or trained animals 24h after LPS application. **(H)** The fraction of labeled cells (relative to ES HSCs) in BM HSPCs of control, trained and untrained *Fgd5*^ZsGreen:CreERT2^/*R26*^LSL-tdRFP^ animals. **(I)** Average number of additional divisions in response to LPS in trained and untrained *R26*^rtTA^/*Col1A1*^H2B-GFP^ animals (dotted line: mean of controls).

## Discussion

Quiescent tip HSCs with high transplantation potential are regarded as a reserve population that directly fuels emergency hematopoiesis (King and Goodell, 2011; Takizawa et al., 2012; Trumpp et al., 2010) We investigated the contribution of these cells to hematopoietic stress responses under native conditions and found that the main flux from HSCs in response to perturbations reached their immediate progeny, but not more distant populations, thereby raising the question of how hematopoietic stress responses are hierarchically organized.

Myeloablative chemotherapy and irradiation recruited quiescent HSCs into cell cycle resulting in significant propagation of HSC label. However, even under these severe, potentially life-threatening conditions, only a minority of HSCs differentiated into their immediate progeny. Of note, even upon transplantation, HSCs alone are insufficient to ensure the short-term survival of the myeloablated host and erythro-myeloid progenitors were shown to be crucial for initial radioprotection until HSCs provide long-term repopulation (Na Nakorn et al., 2002). Hence, HSCs alone are not able to rapidly generate sufficient numbers of mature blood cells even upon myeloablation.

Tip HSCs only marginally contributed to mature blood cell production under conditions mimicking bacterial or viral infection. Neither LPS-induced emergency myelopoiesis nor acute type I IFN signaling robustly induced differentiation of HSCs into committed progenitors or mature hematopoietic cells. Importantly, HSC-derived label did not reach mature myeloid cells faster upon LPS or G-CSF induced myelopoiesis, demonstrating that this stress response was achieved without direct HSC input. While emergency myelopoiesis provoked only minimal cell division of HSPCs on average, type I IFN exposure robustly induced proliferation as previously reported (Bogeska et al., 2022; Essers et al., 2009). Given the massive proliferation of HSPCs in response to escalating doses of pI:C, this protocol induced significant but surprisingly little differentiation of labeled HSCs. We speculate that a massive type I IFN response provoked cell death in HSPCs and proliferation of residual cells compensated for this loss without simultaneously increasing the differentiation flux from HSCs. Indeed, exaggerated type I IFN was shown to induce cell death (Chawla-Sarkar et al., 2003), however, HSCs were reported to be refractory to its destructive effects (Pietras et al., 2014; Wu et al., 2018). Our finding that HSCs participate rarely, if at all, in hematopoietic stress responses contrasts with the frequent observation that severe or chronic infection and inflammation profoundly reduce HSC transplantation potential (reviewed by Caiado et al., 2021). This phenomenon has usually been linked to attrition of HSC during stress responses (Walter et al., 2015). The observation that only a minor fraction of humans ever suffer from bone marrow failure in spite of life-long recurrence of infections also argues for a re-evaluation of this concept.

The main regenerative burden of the hematopoietic system under steady-state conditions is the production of RBCs and Plts (Cosgrove et al., 2021). Interestingly, the rapid recovery after a severe and potentially life-threating reduction of either cell type occurred without significant HSC input. RBCs can solely be replaced by massive proliferation of progenitor cells, while thousands of Plts are generated by a single Mk. Accordingly, a direct differentiation of HSCs into Mks is well documented (Grinenko et al., 2018; Morcos et al., 2022; Rodriguez-Fraticelli et al., 2018; Yamamoto et al., 2013). Both amplification strategies, however, might be too slow for emergency responses emanating from the relatively small HSC population. To rapidly put out 10^9^ RBCs, 10^4^ HSCs would have to undergo simultaneous massive expansion and simultaneous maturation within several days. Similarly, the process of endoreplication in nascent Mks might take too long to boost emergency thrombopoiesis. Reports of an inflammation-induced stem-like MkP population could result from omitting Sca-1 from the HSPC gating strategy (Haas et al., 2015). Thus, these cells could actually correspond to CD48^-/lo^ MkPs, which immuno-phenotypically resemble HSCs except for lacking Sca-1 and EPCR expression.

Innate immune memory is thought to be imprinted in the HSC epigenome (de Laval et al., 2020; Li et al., 2022) and many mature myeloid cells are too short-lived to facilitate long-term alterations of innate immune responses. However, we found a quantitatively unchanged HSC response to LPS stimulation in previously β-Glucan-trained animals, which likely argues against innate immune memory in primitive HSCs. Moreover, our observation that upon inflammatory perturbations, HSC-derived label hardly reaches committed progenitors and even fewer mature blood cells, renders innate immune memory in HSCs less plausible. Given the underappreciated self-renewal capacity and longevity of progenitors under native conditions (Patel et al., 2022), these cells are in principle well-equipped to store information about previous pathogen encounters.

Our results show that tip HSCs do not represent a reserve population in physiological hematopoietic emergencies, such as blood loss and inflammation. Instead, we speculate that HSCs slowly differentiate or “trickle-down” to constantly replenish and rejuvenate progenitor pools at a very low rate, while stress conditions only mildly amplify this process. Consequently, hematopoietic perturbations must be primarily compensated by extensive self-renewal of more differentiated progenitors.

## Limitations of the study

We acknowledge that our *Fgd5*^ZsGreen:CreERT2^/*R26*^LSL-tdRFP^ fate mapping model labels about 10 % of ES HSCs, which then serve as a proxy for the entire tip HSC population. This assumption is substantiated by the following observations: (i) the labeling of ES HSCs and total immuno-phenotypic HSC completely equilibrated starting from one year after induction (Figure 1D), (ii) this label steadily propagated in a linear fashion to all mature blood cell lineages and (iii) most of the long-term transplantation potential is confined to ES HSCs and equal in labeled and unlabeled ES HSCs (Morcos et al., 2022). Our study is further complemented by H2B-GFP proliferation tracking, in which all cells of a given BM population are labeled and traced, but we tracked the average proliferative behavior of a cell population and potentially neglected its heterogeneous cell cycle activity. Finally, we recognize that innate immune training might primarily manifest in qualitative alterations of cell properties (e.g., increased anti-microbial activity of immune cells), and our mouse models would not report such changes.

## Acknowledgement

We thank Christa Haase, Livia Schulze and Luisa Röbisch for expert technical assistance; Triantafyllos Chavakis and Lydia Kalafati for scientific discussions. This work was funded by the German Research Council (DFG, GE3038/1-1 to A.G. and TR237 (B17) to A.R.).

## Author contribution

C.M. performed experiments with help from N.D. and M.C. T.G. provided methodology. A.G. conceived and supervised the study. A.G. and C.M. wrote the paper; A.R. revised the manuscript.

## Star Methods

### Mice

All animal experiments were performed in accordance with institutional guidelines and the German Law for Protection of Animals approved by Landesdirektion Dresden (TVV 91/2017). Mice were housed in individually ventilated cages under specific-pathogen free environment at the Experimental Center of the Medical Faculty, TU Dresden. *R26*^rtTA/rtTA^/*Col1A1*^H2B-GFP/H2B-GFP^ (Foudi et al., 2008) mice were induced with DOX (2g/kg) via chow (Ssniff Spezialdiäten) for 1-2 weeks ad libitum. *Fgd5*^ZsGreen:CreERT2/wt^/*R26*^LSL-tdRFP/LSL-tdRFP^ (Gazit et al., 2014; Luche et al., 2007) mice were induced by oral gavage of TAM (0.2 mg/g body weight (BW)) twice 3-4 days apart. Fate mapping data from *Fgd5*^ZsGreen:CreERT2^/*R26*^LSL-tdRFP^ mice shown in Figures 1C and D, and 5-FU perturbation data shown in Figures 1G-P and S2C-H was previously published (Morcos et al., 2022). 5-FU (150 μg/g BW, Applichem) was administered via intravenous (i.v.) injection. Whole body irradiation was performed using a Yxlon Maxi Shot X-ray tube at a dose of 2 Gy. Recombinant human G-CSF (Filgrastim) (0.3 μg/g BW, Neupogen, Amgen) was administered by subcutaneous (s.c.) injection on 5 consecutive days. pI:C (5 μg/g BW, Poly(I:C), Invivogen) was administered via intraperitoneal injection (i.p.) either one (“single” protocol) or 8 times (“repetitive” protocol, 2 injections/week for 4 consecutive weeks. LPS (from E. coli O111:B4, Invivogen, 35 μg/mouse in two out of four experiments and adjusted to 1.4 μg/g BW in the remaining experiments) and β-glucan peptide (from Trametes versicolor, Invivogen, 1mg/mouse in two out of four experiments and adjusted to 0.04 mg/g BW in the remaining experiments) were injected intraperitoneal. Rabbit anti-mouse thrombocyte serum (150 μl/mouse, WAK-Chemie Medical GmbH) or mock serum were administered i.p. either once (“acute” protocol) or seven times (“chronic” protocol, injection every other day).

### Cell Preparation

Extraction of whole BM cells was achieved by crushing long bones with mortar and pestle using PBS/2% FCS/2 mM EDTA and filtering through a 100 μm mesh and a secondary filtering step through a 30 μm mesh after erythrocyte lysis in hypotonic NH_4_Cl-buffer. Hematopoietic lineage^+^ cells were depleted from samples prior to final staining with the lineage cell depletion kit (Miltenyi Biotec).

Peripheral blood was drawn into glass capillaries by retro-bulbar puncture directly into EDTA-coated tubes (Sarstedt). For identification of RFP^+^ platelets and erythrocytes, 1-2 μl of whole blood was mixed with PBS/2%FCS/2mM EDTA and incubated with monoclonal antibodies. For leukocyte analysis, erythrocyte lysis in hypotonic NH_4_Cl-buffer was performed twice for 5 min before cells were stained with monoclonal antibodies (for gating of PB populations see Figure S1B, C). For hemograms, blood was diluted 1:5 in isotonic NaCl solution and analyzed on an XT-2000i Vet analyzer (Sysmex).

Spleen cell suspensions were prepared by mashing spleens using PBS/2% FCS/2 mM EDTA through a 100 μm mesh, subsequent erythrocyte lysis in hypotonic NH_4_Cl-buffer and antibody staining.

Peritoneal cells were obtained by flushing the peritoneal cavity with 4ml PBS/2% FCS/2 mM EDTA and filtering the aspirate through a 100 μm mesh (for gating of peritoneal cells see Figure S1D).

### Flow cytometry

Cell suspensions from BM, spleen, peritoneal lavage or PB samples were stained with antibodies (see Key Rescource Table) in PBS/2% FCS/2mM EDTA for 30-40 min, washed twice and analyzed on either FACS Canto, ARIA II SORP, Aria II, ARIA III (all from BD Biosciences, Heidelberg, Germany) or MACSquant (Miltenyi) flow cytometers. FlowJo V9.9 and V10 software (Tree Star) was used for data analysis, and gates were set using Fluorescence-Minus-One controls. Details on gating strategies are summarized in Figure S1A-D. Absolute numbers of lin^-^ BM cells were determined employing a Miltenyi MACSquant flow cytometer.

### Plasma cytokine concentration

PB samples were centrifuged for 10 min at 1000 g within 30 minutes of blood collection. Plasma supernatants were collected and frozen at −20°C until analysis. For measuring plasma cytokine concentration, we used the LEGENDplex™ assay kit (13 plex mouse inflammation panel, BioLegend) according to the manufacturer’s instructions. Flow cytometric analysis was performed on a MACSquant flow cytometer, data analysis was performed via the online available LEGENDplex™ Data Analysis Software Suite.

### Data normalization and statistics

*RFP label propagation data* from various BM and PB populations of induced *Fgd5*^ZsGreen:CreERT2^/*R26*^LSL-tdRFP^ mice was subjected to the following normalization steps:

1. Normalization to ES HSCs, which are the ultimate source of label propagation in *Fgd5*^ZsGreen:CreERT2^/*R26*^LSL-tdRFP^ mice (as explained in Figure 1B-D); the measured label in any given population x of animal 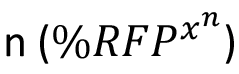 was divided by the measured label in ES HSCs of the same animal 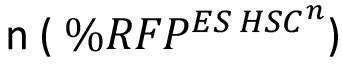 to receive the fraction of label in ES HSC (*Label_norm_*:

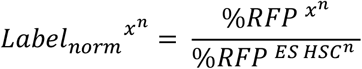
2. The percentage of label increase in a certain population x caused by the stress response was calculated by subtracting the normalized arithmetic mean label in control animals 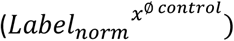 from the normalized label in a treated specimen n (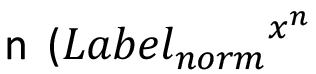, further explained in Figure S2F):

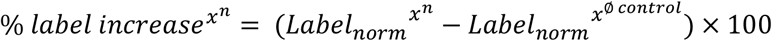

For clarity, the normalized RFP label propagation data was summarized in radar charts (Figures 1O,2P, 3M). To this end, mean label increase data from perturbed animals were used (colored lines). The % of label increase in controls averages at zero within each individual experiment (dotted grey lines). To obtain an estimate of natural fluctuation, the % of label increase in control-treated animals from all perturbation experiments (n=41, 8 experiments) were collected and the standard deviation was calculated (grey shaded areas, see also Figure S2G-H). Radar charts were plotted using the *radarchart* function of the *fsmb* R-package.

*H2B-GFP label retention data* from *R26*^rtTA^/*Col1A1*^H2B-GFP^ mice was normalized in the following way:

As a result of perturbation, the mean H2B-GFP label in population x of animal 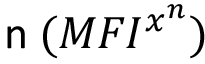 may dilute faster than in the average of all control animals 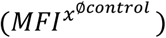; hence, the number of possible bisections of fluorescence intensity values between a treated animal and the control group average corresponds to the number of additional divisions (further explained in Figure S2D-E). Depending on the sex of animal n, average values from either male or female control animals were used to perform the calculation, since initial H2B-GFP fluorescent levels after DOX induction differ between male and female animals (Morcos et al., 2020).

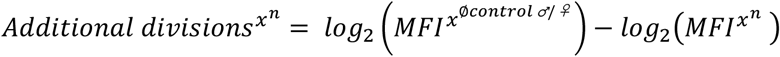

### Display and statistics

All graphs shown in main or supplementary figures are designed with the following properties: Individual mice are shown, box-plots show medians, lower and upper quartiles; dotted lines represent means of untreated controls; bar graphs show means; significance between perturbed and control mice was calculated by an unpaired Student’s t-test except for Figures 1E-G, S2B, S3L, S4M and O, were a 1way ANOVA with Tukey post test was applied (* p ≥ 0.05; ** p ≥ 0.01, *** p ≥ 0.001, non-significant differences are not indicated.)

---

**Figure S1.**
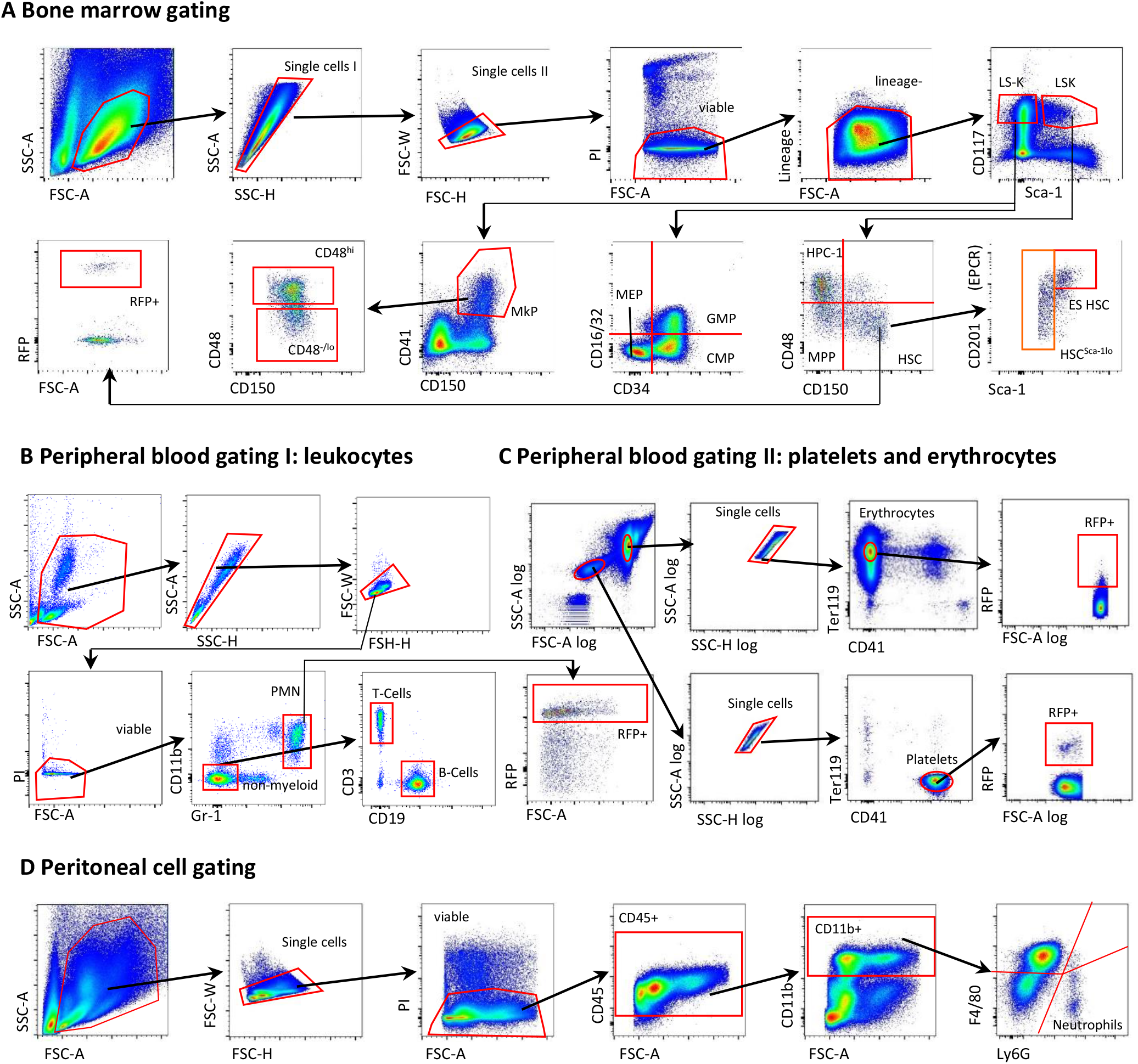
related to Figure 1: Flow cytometry gating strategies. **(A)** Gating of lineage-negative (lin^-^) BM to identify hematopoietic stem and progenitor populations; (*LS^-^K*, lin^-^ Sca-1^-^ CD117^+^ cells; *LSK*, lin^-^ Sca-1^+^ CD117^+^ cells; *megakaryocyte progenitor (MkP*), LS^-^K CD41^+^ CD150^+^; *common myeloid progenitor (CMP*), LS^-^K CD16/32^-^ CD34^+^; *restricted hematopoietic progenitor 1 (HPC-1*), LSK CD48^hi^ CD150^-^; *restricted hematopoietic progenitor 2 (HPC-2*), LSK CD48^hi^ CD150^+^; *multipotent progenitor (MPP*), LSK CD48^-/lo^ CD150^-^; *hematopoietic stem cell (HSC*), LSK CD48^-/lo^ CD150^+^; *ES HSC*, CD201^hi^ Sca-1^hi^ LSK CD48^-/lo^ CD150^+^; *Sca-1^lo^ HSC*, Sca-1^lo^ LSK CD48^-/lo^ CD150^+^;). The RFP-labeling frequencies in BM populations of induced *Fgd5*^ZsGreen:CreERT2^*/R26*^LSL-tdRFP^ mice were analyzed using two different staining panels (panel 1: HSC, MPP, HPC-1 and MkP, panel 2: CMP, GMP, MEP). **(B)** Gating of peripheral blood (PB) leukocytes after erythrocyte lysis (*polymorphonuclear leukocytes* (*PMN*, granulocytes), CD11b^+^ Gr-1^+^; *B-Cells*, CD11b^-^ Gr-1^-^ CD19^+^; *T-Cells*, CD11b^-^ Gr-1^-^ CD3^+^). Representative gating of RFP^+^ neutrophils is shown. **(C)** Identification of RFP^+^ erythrocytes (CD41^-^ Ter119^+^) and platelets (CD41^+^ Ter119^-^) in PB. **(D)** Representative gating of peritoneal cavity (PC) neutrophils (CD45^+^CD11b^+^ Ly6-G^+^F4/80^-/lo^).

**Figure S2.**
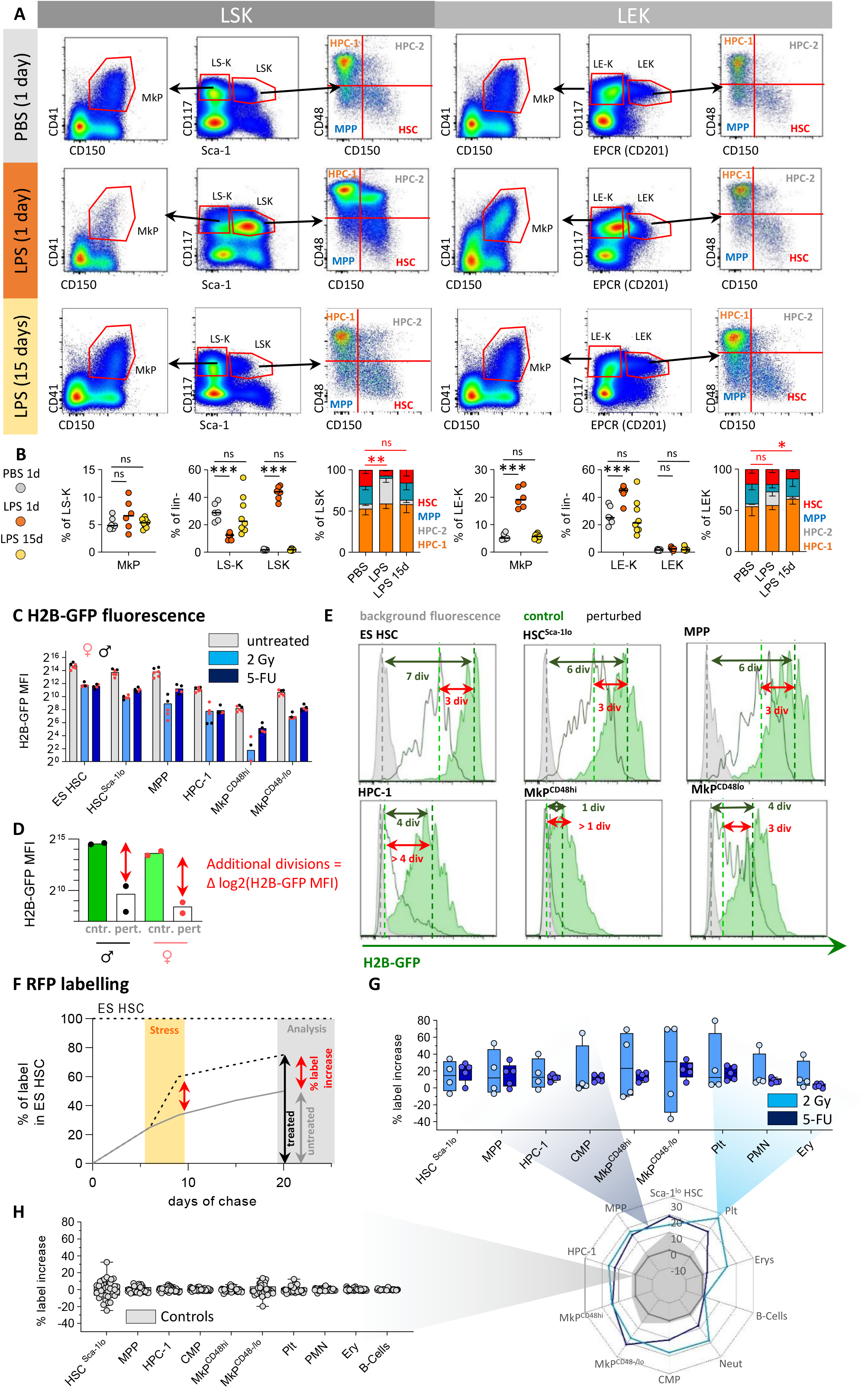
related to Figure 1: HSPC immuno-phenotypes during inflammation; transformation of label propagation and proliferation tracking data. **(A-B)** Under conditions of acute inflammation, canonical immuno-phenotypes of BM stem and progenitor cell populations become unreliable. Most prominently, Sca-1, a marker used to discriminate early HSPCs from committed progenitors, is upregulated by proinflammatory cytokines including LPS (**A**, compare upper and middle row of dot plots, 1 day after PBS or LPS treatment, respectively), which leads to massive alterations in identity and abundance of BM populations (**B**, row of data plots) and exemplifies that comparison of data acquired under steady-state and inflammatory conditions is implausible. An alternative gating strategy (Vazquez et al., 2015) using EPCR (CD201) as a replacement for Sca-1 (**A**, LEK-gating, right dot plots) appears more reliable upon acute inflammation (**B**). Both *Fgd5*^ZsGreen:CreERT2^*/R26*^LSL-tdRFP^ and *R26*^rtTA^/*Col1A1*^H2B-GFP^ reporter mouse models cumulatively record label propagation and cell division, respectively, and were read-out after the perturbations had settled (≥ 2weeks) and faithful marker expression was restored (A, lower row of dot plots, 14 days after LPS). All analyses of BM HSPCs performed during acute perturbations (within 48 h after perturbation) in this study employed the more reliable LEK gating strategy (Figures 2G, H and S3I-L) **(C)** *R26*^rtTA^/*Col1A1*^H2B-GFP^ animals (same mice as in Figure 1N-O) were Dox-pulsed and exposed to either 2 Gy (n=5), 5-FU (n=5) or left untreated (n=6). H2B-GFP mean fluorescence intensity (MFI) values of BM populations of myeloablated and control animals are shown (Black dots = males, red dots = females). **(D)** Male and female mice show differences in maximum H2B-GFP MFI values directly after induction. Thus, additional divisions (Δ log2(H2B-GFP MFI)) were calculated separately for male (black) and female (red) mice by subtracting the H2B-GFP value in a certain population of a treated animal from average H2B-GFP values found in the corresponding population of either male or female control animals. Graph shows gender-sorted H2B-GFP MFI values on ES HSCs from control (green bars) or 8x pI:C-treated (open bars) animals. **(E)** The average number of additional cell divisions in response to perturbation was estimated by H2B-GFP proliferation tracking. The difference of mean H2B-GFP intensities (MFI) between perturbed and control was calculated (red arrows, Δ log2(H2B-GFP MFI)). Histograms show representative examples of BM populations isolated from pulsed and chased *R26*^rtTA^/*Col1A1*^H2B-GFP^ animals perturbed with 5-FU (light green, open histograms “perturbed”), or left untreated (dark green, filled histograms “control”). Leaky background H2B-GFP fluorescence was estimated from an untreated, uninduced *R26*^rtTA^/*Col1A1*^H2B-GFP^ animal (grey histogram, “background”). The maximum number of traceable additional divisions (green arrows) varies between BM populations and depends on the mean distance between H2B-GFP fluorescent intensity in unperturbed control and background animals. When the H2B-GFP fluorescence distribution of a population approaches the range of background controls, further proliferation is not faithfully trackable, since division no longer results in halving of H2B-GFP (see Morcos et al. (2020) for further discussion). For example, the near complete dilution of H2B-GFP in progenitors responding to 5-FU (Figures S2 C, E in HPC-1 and MkP populations and Figure 1O), ionizing radiation (Figures S2C and 1O), or repetitive pI:C (Figures 2O and S3T) perturbation indicates exhaustion of the mitotic tracker. Moreover, differences in the number of additional division between distinct BM populations exposed to the same perturbation should be treated with caution as upon vigorous proliferation and accordingly strong H2B-GFP dilution, these differences might reflect merely the varying number of traceable additional divisions in each population (e.g. up to ^~^6-7 and ^~^3-4 additional divisions traceable in HSCs and HPC-1s, respectively). However, this caveat does not apply to the comparison of the same HSPC population between perturbed and un-perturbed controls animals (i.e. the additional divisions themselves, which are faithfully trackable until the H2B-GFP reaches background levels). Mitotic history in response to perturbation is challenging to estimate in late hematopoietic progenitors, which rapidly proliferate already in steady state and therefore quickly dilute H2B-GFP to background level in unperturbed controls. **(F)** RFP label propagation from initially labeled ES HSC in TAM-induced *Fgd5*^ZsGreen:CreERT2^*/R26*^LSL-tdRFP^ is a slow, but continuous process (grey curve, fictional population). During acute stress responses (yellow shaded area) this population accelerates label acquisition, given that the source of the label increases their output (black, dotted curve). Upon recovery from hematopoietic perturbation, the differentiation- and proliferation rates revert back to steady state. The stress-induced differences in population labeling between treated and untreated animals are preserved and can be recovered at later time points, when reliable HSPC surface marker expression is re-gained. To extract the net effect of each perturbation on RFP label propagation, we subjected the data to the following transformation steps: (i) normalization to the source (i.e. normalization of label (% of RFP^+^ cells) of any given population to the labeling frequency present in the same animal’s ES HSC population, and (ii) calculating the offset between normalized data from treated (black arrow) and untreated (grey arrow) animals, which yields the % of label increase (red arrow, net effect) caused by the perturbation. **(G-H)** *Fgd5*^ZsGreen:CreERT2^*/R26*^LSL-tdRFP^ mice were TAM-induced and exposed to 2 Gy γ-irradiation (n=4), 5-FU (n=5) or left untreated (n=4) and BM and PB were analyzed for RFP expression (same animals as in Figures 1 G-M). **(G)** Transformed labeling data revealing the net label increase (% relative to ES HSC label) in BM population of myeloablated animals after subtraction of the mean labeling of unperturbed control mice (transformation was performed as described in Figure S2F). **(H)** Previously TAM-induced *Fgd5*^ZsGreen:CreERT2^*/R26*^LSL-tdRFP^ mice (controls) from all experiments (n=41, 8 independent experiments), that were left unperturbed. Means are defined as zero and serve as a reference point for perturbation effects throughout the study. Variance (SD) of control-treated animals is shown in radar plots in Figures 1P, 2P and 3M as shaded areas.

**Figure S3.**
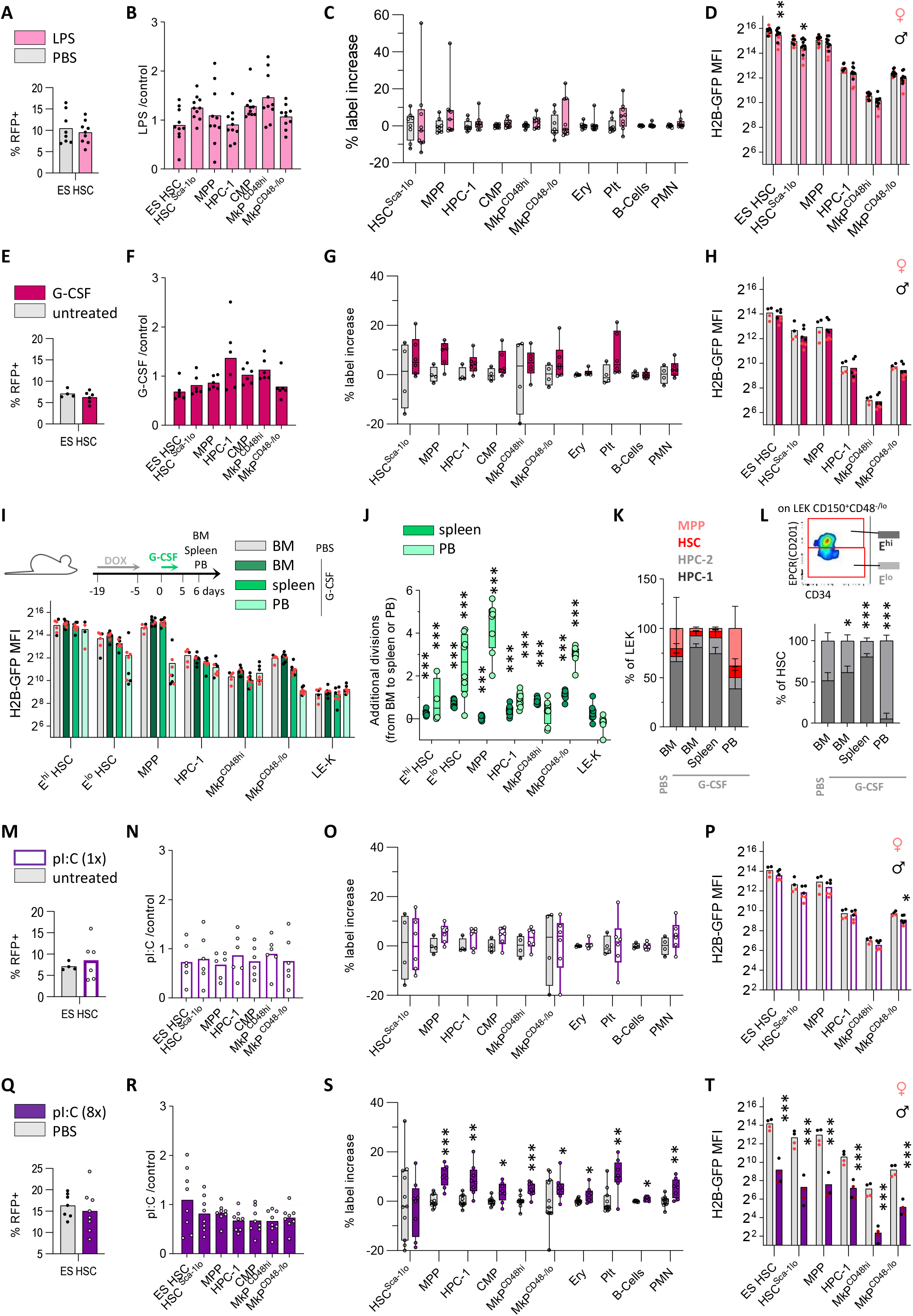
related to Figure 2: Fate mapping and proliferation tracking in mice exposed to inflammatory cytokine signaling. **(A-C)** *Fgd5*^ZsGreen:CreERT2^*/R26*^LSL-tdRFP^ mice were TAM-induced and i.p. injected with LPS (n=8) or PBS (n=12) and BM and PB were analyzed for RFP expression (same animals as in Figures 2A-D; data pooled from two independent experiments). **(A)** RFP labeling in ES HSCs. **(B)** Ratios of relative BM compartment sizes (% cells among total lin-BM cells) between perturbed and control (dotted line, individual control mice not shown) animals. **(C)** Net label increase in LPS-perturbed animals after subtraction of label from saline-injected control animals. Resulting means of BM populations were plotted in the radar chart shown in Figure 2P). **(D)** Previously DOX-induced *R26*^rtTA^/*Col1A1*^H2B-GFP^ animals (same mice as in Figure 2E; data pooled from two independent experiments) were injected with LPS (n=14) or PBS (n=13). H2B-GFP MFI values of BM populations of perturbed and control animals are shown (Black dots = males, red dots = females). **(E-H)** *Fgd5*^ZsGreen:CreERT2^*/R26*^LSL-tdRFP^ (**E-G,** n=6-8/condition) and *R26*^rtTA^/*Col1A1*^H2B-GFP^ (**H,** n=4-7/condition) mice were s.c. injected with G-CSF or PBS (same animals as in Figure 2F-J; display of data as in Figure S3A-D). **(E)** RFP labeling in ES HSCs; **(F)** BM compartment sizes; **(G)** net label increase upon perturbation (see also Figure 2P) and **(H)** H2B-GFP MFIs are shown. **(I-L)** *R26*^rtTA^/*Col1A1*^H2B-GFP^ animals were s.c. injected with G-CSF (n=8) or PBS (n=6) for 5 consecutive days and HSPCs (LEK-gated, see Figure S2A) were analyzed one day after the last injection. **(I)** H2B-GFP MFI values of HSPC populations in BM, spleen or PB (**J**). The average additional divisions of HSPCs mobilized to spleen or PB in comparison to BM was calculated within each G-CSF injected individual. **(K)** Composition of the LEK compartment subdivided by CD48 and CD150 expression. **(L)** Composition of the HSC compartment (LEK CD48^-/lo^CD150^+^) subdivided by CD201 (EPCR) (dot plot). **(M-P)** *Fgd5*^ZsGreen:CreERT2^/*R26*^LSL-tdRFP^ (**M-O,** n=6-7/condition) and *R26*^rtTA^/*Col1A1*^H2B-GFP^ (**P,** n=4-6/condition) mice were injected once with p:IC or PBS (same animals as in Figures 2K-O; display of data as in Figure S3A-D). **(M)** RFP label in ES HSCs; **(N)** BM compartment sizes; **(O)** net label increase upon treatment (see also Figure 2P) and **(P)** H2B-GFP MFIs are shown. **(Q-T)** *Fgd5*^ZsGreen:CreERT2^/*R26*^LSL-tdRFP^ (**Q-S,** n=7-8/condition) and *R26*^rtTA^/*Col1A1*^H2B-GFP^ (**T,** n=4/condition) mice were injected 8x with pI:C or PBS (same animals as in Figures 2K-O; display of data as in Figure S3A-D). **(Q)** RFP label in ES HSCs; **(R)** BM compartment sizes; **(S)** net label increase upon treatment (see also Figure 2P) and **(T)** H2B-GFP MFIs are shown.

**Figure S4.**
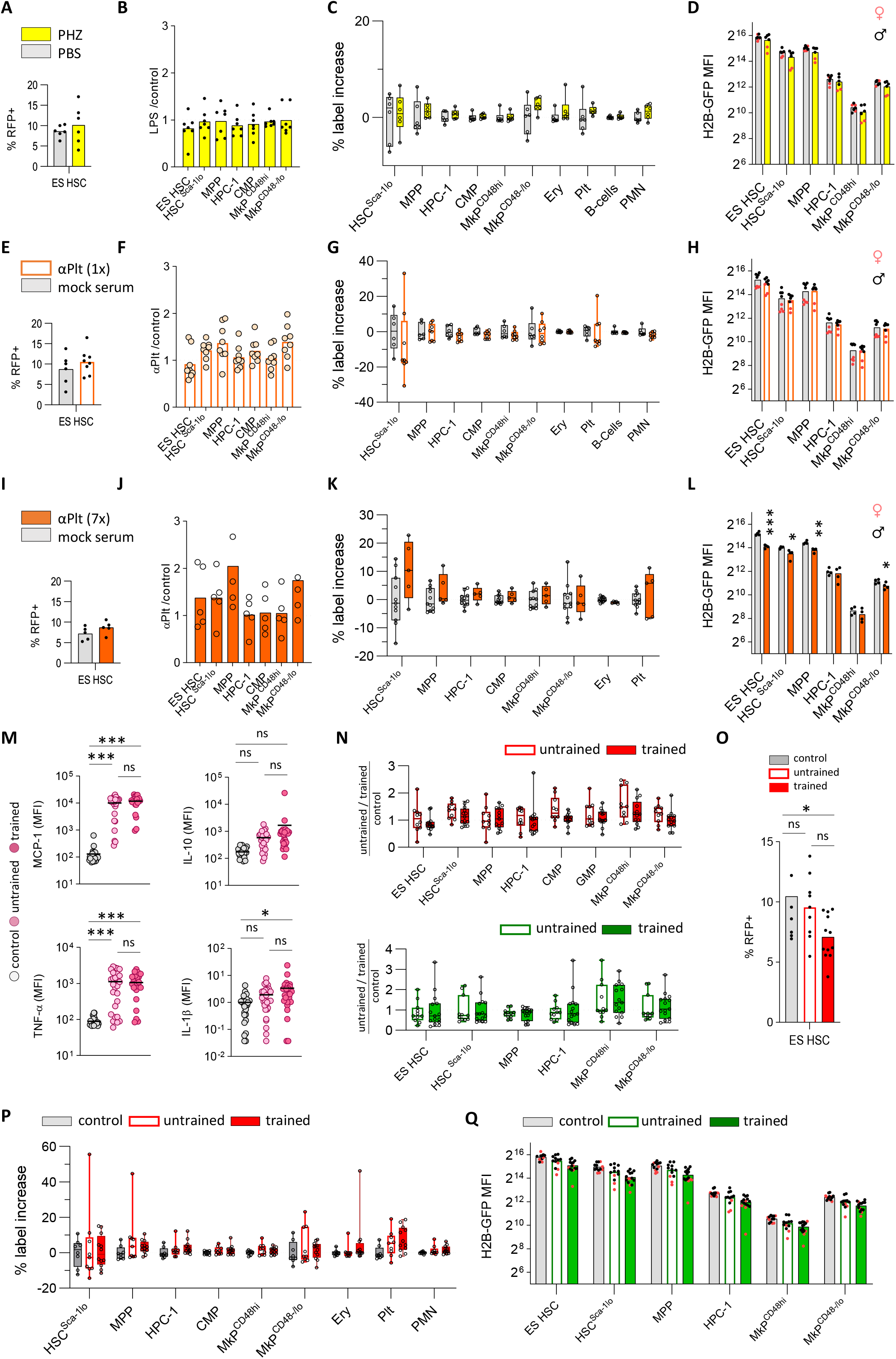
related to Figures 3 and 4: Fate mapping and proliferation tracking in mice exposed to blood cell loss and innate immune training. **(A-C)** *Fgd5*^ZsGreen:CreERT2^/*R26*^LSL-tdRFP^ mice were TAM-induced and i.p. injected with PHZ (n=7) or PBS (n=6) and BM and PB were analyzed for RFP expression (same animals as in Figure 3A-D). **(A)** RFP labeling in ES HSCs; **(B)** ratios of relative BM compartment sizes (% cells among total lin-BM cells) between perturbed and control (dotted line, individual control mice not shown) animals and **(C)** net label increase in LPS-perturbed animals after subtraction of label from saline-injected control animals. Resulting meanof BM populations were plotted in the radar chart shown in Figure 3M). **(D)** Previously DOX-induced *R26*^rtTA^/*Col1A1*^H2B-GFP^ animals (same mice as in Figure 3E) were i.p. injected with PHZ (n=6) or PBS (n=7). H2B-GFP MFI values of BM populations of PHZ-injected and control animals are shown (Black dots = males, red dots = females). **(E-H)** *Fgd5*^ZsGreen:CreERT2^/*R26*^LSL-tdRFP^ (**E-G,** n=6-8/condition) and *R26*^rtTA^/*Col1A1*^H2B-GFP^ (**H,** n=8/condition) mice were injected once with αPlt-serum or mock serum (same animals as in Figure 3F-L). **(E)** RFP labeling in ES HSCs; **(F)** BM compartment sizes; **(G)** net label increase upon treatment (see also Figure 3M) and **(H)** H2B-GFP MFIs are shown. **(I-L)** *Fgd5*^ZsGreen:CreERT2^/*R26*^LSL-tdRFP^ (**I-K,** n=5/condition) and *R26*^rtTA^/*Col1A1*^H2B-GFP^ (**L,** n=4/condition) mice were i.p. injected with αPlt-serum or mock serum repeatedly (7 injections within 12 days, same animals as in Figure 3F-L). **(I)** RFP labeling in ES HSCs; **(J)** BM compartment sizes; **(K)** net label increase upon treatment (see also Figure 3M) and **(L)** H2B-GFP MFIs are shown. **(M-Q)** Innate immune training was stimulated in *Fgd5*^ZsGreen:CreERT2^/*R26*^LSL-tdRFP^ (red, n=8-12/condition, 2 independent experiments) and *R26*^rtTA^/*Col1A1*^H2B-GFP^ (green, n=13-14/condition, 2 independent experiments) animals (same mice as in Figure 4), **(M) L**evels of MCP-1, IL-10, tumor necrosis factor (TNF) and IL-1β in plasma of control, untrained and trained animals 24 h after PBS or LPS application. **(N)** Ratios of relative compartment sizes (% cells among lin-BM cells) of trained and untrained *Fgd5*^ZsGreen:CreERT2^*/R26*^LSL-tdRFP^ (upper plot) and *R26*^rtTA^/*Col1A1*^H2B-GFP^ (lower plot) animals compared to controls (dotted line, individual animals not shown). **(O)** RFP labeling in ES HSCs; **(P)** net label increase in untrained and trained animals after subtraction of label from PBS-injected control animals. **(Q)** H2B-GFP MFI values of BM populations isolated from trained, untrained and control *R26*^rtTA^/*Col1A1*^H2B-GFP^ animals.

## References

Bernitz, J.M., Daniel, M.G., Fstkchyan, Y.S., and Moore, K. (2017). Granulocyte colony-stimulating factor mobilizes dormant hematopoietic stem cells without proliferation in mice. Blood 129, 1901–1912.

Beutler, B.A. (2009). TLRs and innate immunity. Blood 113, 1399–1407.

Boettcher, S., Gerosa, R.C., Radpour, R., Bauer, J., Ampenberger, F., Heikenwalder, M., Kopf, M., and Manz, M.G. (2014). Endothelial cells translate pathogen signals into G-CSF–driven emergency granulopoiesis. Blood 124, 1393–1403.

Bogeska, R., Mikecin, A.-M., Kaschutnig, P., Fawaz, M., Büchler-Schäff, M., Le, D., Ganuza, M., Vollmer, A., Paffenholz, S.V., Asada, N., et al. (2022). Inflammatory exposure drives long-lived impairment of hematopoietic stem cell self-renewal activity and accelerated aging. Cell Stem Cell 29, 1273–1284.e1278.

Bowling, S., Sritharan, D., Osorio, F.G., Nguyen, M., Cheung, P., Rodriguez-Fraticelli, A., Patel, S., Yuan, W.C., Fujiwara, Y., Li, B.E., et al. (2020). An Engineered CRISPR-Cas9 Mouse Line for Simultaneous Readout of Lineage Histories and Gene Expression Profiles in Single Cells. Cell 181, 1410–1422.e1427.

Bujanover, N., Goldstein, O., Greenshpan, Y., Turgeman, H., Klainberger, A., Scharff, Y.e., and Gazit, R. (2018). Identification of immune-activated hematopoietic stem cells. Leukemia 32, 2016–2020.

Busch, K., Klapproth, K., Barile, M., Flossdorf, M., Holland-Letz, T., Schlenner, S.M., Reth, M., Höfer, T., and Rodewald, H.-R. (2015). Fundamental properties of unperturbed haematopoiesis from stem cells in vivo. Nature 518, 542–546.

Caiado, F., Pietras, E.M., and Manz, M.G. (2021). Inflammation as a regulator of hematopoietic stem cell function in disease, aging, and clonal selection. Journal of Experimental Medicine 218.

Chawla-Sarkar, M., Lindner, D.J., Liu, Y.F., Williams, B.R., Sen, G.C., Silverman, R.H., and Borden, E.C. (2003). Apoptosis and interferons: Role of interferon-stimulated genes as mediators of apoptosis. Apoptosis 8, 237–249.

Cosgrove, J., Hustin, L.S.P., de Boer, R.J., and Perié, L. (2021). Hematopoiesis in numbers. Trends in Immunology 42, 1100–1112.

de Laval, B., Maurizio, J., Kandalla, P.K., Brisou, G., Simonnet, L., Huber, C., Gimenez, G., Matcovitch-Natan, O., Reinhardt, S., David, E., et al. (2020). C/EBPβ-Dependent Epigenetic Memory Induces Trained Immunity in Hematopoietic Stem Cells. Cell Stem Cell 26, 657–674.e658.

Demerdash, Y., Kain, B., Essers, M.A.G., and King, K.Y. (2021). Yin and Yang: The dual effects of interferons on hematopoiesis. Experimental Hematology 96, 1–12.

Essers, M.A.G., Offner, S., Blanco-Bose, W.E., Waibler, Z., Kalinke, U., Duchosal, M.A., and Trumpp, A. (2009). IFNalpha activates dormant haematopoietic stem cells in vivo. Nature 458, 904–908.

Foudi, A., Hochedlinger, K., Van Buren, D., Schindler, J.W., Jaenisch, R., Carey, V., and Hock, H. (2008). Analysis of histone 2B-GFP retention reveals slowly cycling hematopoietic stem cells. Nature Biotechnology 27, 84–90.

Gazit, R., Mandal, P.K., Ebina, W., Ben-Zvi, A., Nombela-Arrieta, C., Silberstein, L.E., and Rossi, D.J. (2014). Fgd5 identifies hematopoietic stem cells in the murine bone marrow. Journal of Experimental Medicine 211, 1315–1331.

Greenbaum, A.M., and Link, D.C. (2011). Mechanisms of G-CSF-mediated hematopoietic stem and progenitor mobilization. Leukemia 25, 211–217.

Grinenko, T., Eugster, A., Thielecke, L., Ramasz, B., Krüger, A., Dietz, S., Glauche, I., Gerbaulet, A., von Bonin, M., Basak, O., et al. (2018). Hematopoietic stem cells can differentiate into restricted myeloid progenitors before cell division in mice. Nature Communications 9, 1898.

Haas, S., Hansson, J., Klimmeck, D., Loeffler, D., Velten, L., Uckelmann, H., Wurzer, S., Prendergast, Aine M., Schnell, A., Hexel, K., et al. (2015). Inflammation-Induced Emergency Megakaryopoiesis Driven by Hematopoietic Stem Cell-like Megakaryocyte Progenitors. Cell Stem Cell 17, 422–434.

Kanayama, M., Izumi, Y., Yamauchi, Y., Kuroda, S., Shin, T., Ishikawa, S., Sato, T., Kajita, M., and Ohteki, T. (2020). CD86-based analysis enables observation of bona fide hematopoietic responses. Blood 136, 1144–1154.

Kanda, T., Sullivan, K.F., and Wahl, G.M. (1998). Histone-GFP fusion protein enables sensitive analysis of chromosome dynamics in living mammalian cells. Current Biology 8, 377–385.

King, K.Y., and Goodell, M.A. (2011). Inflammatory modulation of HSCs: viewing the HSC as a foundation for the immune response. Nature Reviews Immunology 11, 685–692.

Li, X., Wang, H., Yu, X., Saha, G., Kalafati, L., Ioannidis, C., Mitroulis, I., Netea, M.G., Chavakis, T., and Hajishengallis, G. (2022). Maladaptive innate immune training of myelopoiesis links inflammatory comorbidities. Cell 185, 1709–1727.e1718.

Luche, H., Weber, O., Nageswara Rao, T., Blum, C., and Fehling, H.J. (2007). Faithful activation of an extra bright red fluorescent protein in “knock in” Cre reporter mice ideally suited for lineage tracing studies. Eur J Immunol 37, 43–53.

Mitroulis, I., Ruppova, K., Wang, B., Chen, L.-S., Grzybek, M., Grinenko, T., Eugster, A., Troullinaki, M., Palladini, A., Kourtzelis, I., et al. (2018). Modulation of Myelopoiesis Progenitors Is an Integral Component of Trained Immunity. Cell 172, 147–161.e112.

Morcos, M.N.F., Li, C., Munz, C.M., Greco, A., Dressel, N., Reinhardt, S., Sameith, K., Dahl, A., Becker, N.B., Roers, A., et al. (2022). Fate mapping of hematopoietic stem cells reveals two pathways of native thrombopoiesis. Nature Communications 13, 4504.

Morcos, M.N.F., Schoedel, K.B., Hoppe, A., Behrendt, R., Basak, O., Clevers, H.C., Roers, A., and Gerbaulet, A. (2017). SCA-1 Expression Level Identifies Quiescent Hematopoietic Stem and Progenitor Cells. Stem Cell Reports 8, 1472–1478.

Morcos, M.N.F., Zerjatke, T., Glauche, I., Munz, C.M., Ge, Y., Petzold, A., Reinhardt, S., Dahl, A., Anstee, N.S., Bogeska, R., et al. (2020). Continuous mitotic activity of primitive hematopoietic stem cells in adult mice. Journal of Experimental Medicine 217.

Morrison, S.J., Wright, D.E., and Weissman, I.L. (1997). Cyclophosphamide/granulocyte colony-stimulating factor induces hematopoietic stem cells to proliferate prior to mobilization. Proc Natl Acad Sci USA 94, 1908–1913.

Na Nakorn, T., Traver, D., Weissman, I.L., and Akashi, K. (2002). Myeloerythroid-restricted progenitors are sufficient to confer radioprotection and provide the majority of day 8 CFU-S. The Journal of Clinical Investigation 109, 1579–1585.

Naik, S.H., Perié, L., Swart, E., Gerlach, C., van Rooij, N., de Boer, R.J., and Schumacher, T.N. (2013). Diverse and heritable lineage imprinting of early haematopoietic progenitors. Nature 496, 229–232.

Netea, M.G., Domínguez-Andrés, J., Barreiro, L.B., Chavakis, T., Divangahi, M., Fuchs, E., Joosten, L.A.B., van der Meer, J.W.M., Mhlanga, M.M., Mulder, W.J.M., et al. (2020). Defining trained immunity and its role in health and disease. Nature Reviews Immunology 20, 375–388.

Patel, S.H., Christodoulou, C., Weinreb, C., Yu, Q., da Rocha, E.L., Pepe-Mooney, B.J., Bowling, S., Li, L., Osorio, F.G., Daley, G.Q., et al. (2022). Lifelong multilineage contribution by embryonic-born blood progenitors. Nature 606, 747–753.

Pei, W., Feyerabend, T.B., Rössler, J., Wang, X., Postrach, D., Busch, K., Rode, I., Klapproth, K., Dietlein, N., Quedenau, C., et al. (2017). Polylox barcoding reveals haematopoietic stem cell fates realized in vivo. Nature 548, 456–460.

Pietras, E.M., Lakshminarasimhan, R., Techner, J.-M., Fong, S., Flach, J., Binnewies, M., and Passegué, E. (2014). Re-entry into quiescence protects hematopoietic stem cells from the killing effect of chronic exposure to type I interferons. Journal of Experimental Medicine 211, 245–262.

Purton, L.E., and Scadden, D.T. (2007). Limiting Factors in Murine Hematopoietic Stem Cell Assays. Cell Stem Cell 1, 263–270.

Ramasz, B., Krüger, A., Reinhardt, J., Sinha, A., Gerlach, M., Gerbaulet, A., Reinhardt, S., Dahl, A., Chavakis, T., Wielockx, B., et al. (2019). Hematopoietic stem cell response to acute thrombocytopenia requires signaling through distinct receptor tyrosine kinases. Blood 134, 1046–1058.

Rodriguez-Fraticelli, A.E., Weinreb, C., Wang, S.-W., Migueles, R.P., Jankovic, M., Usart, M., Klein, A.M., Lowell, S., and Camargo, F.D. (2020). Single-cell lineage tracing unveils a role for TCF15 in haematopoiesis. Nature 583, 585–589.

Rodriguez-Fraticelli, A.E., Wolock, S.L., Weinreb, C.S., Panero, R., Patel, S.H., Jankovic, M., Sun, J., Calogero, R.A., Klein, A.M., and Camargo, F.D. (2018). Clonal analysis of lineage fate in native haematopoiesis. Nature 553, 212.

Schoedel, K., Morcos, M., Zerjatke, T., Roeder, I., Grinenko, T., Voehringer, D., Göthert, J., Waskow, C., Roers, A., and Gerbaulet, A. (2016). The bulk of the hematopoietic stem cell population is dispensable for murine steady-state and stress hematopoiesis. Blood 128, 2285–2296.

Sender, R., and Milo, R. (2021). The distribution of cellular turnover in the human body. Nature Medicine 27, 45–48.

Sheikh, B.N., Yang, Y., Schreuder, J., Nilsson, S.K., Bilardi, R., Carotta, S., McRae, H.M., Metcalf, D., Voss, A.K., and Thomas, T. (2016). MOZ (KAT6A) is essential for the maintenance of classically defined adult hematopoietic stem cells. Blood 128, 2307–2318.

Sun, J., Ramos, A., Chapman, B., Johnnidis, J.B., Le, L., Ho, Y.-J., Klein, A., Hofmann, O., and Camargo, F.D. (2014). Clonal dynamics of native haematopoiesis. Nature 514, 322–327.

Takahashi, M., Barile, M., Chapple, R.H., Tseng, Y.J., Nakada, D., Busch, K., Fanti, A.K., Säwén, P., Bryder, D., Höfer, T., et al. (2021). Reconciling Flux Experiments for Quantitative Modeling of Normal and Malignant Hematopoietic Stem/Progenitor Dynamics. Stem Cell Reports 16, 741–753.

Takizawa, H., Boettcher, S., and Manz, M.G. (2012). Demand-adapted regulation of early hematopoiesis in infection and inflammation. Blood 119, 2991–3002.

Thomas, E.D., Lochte, H.L., Lu, W.C., and Ferrebee, J.W. (1957). Intravenous Infusion of Bone Marrow in Patients Receiving Radiation and Chemotherapy. New England Journal of Medicine 257, 491–496.

Trumpp, A., Essers, M., and Wilson, A. (2010). Awakening dormant haematopoietic stem cells. Nature Reviews Immunology 10, 201–209.

Vazquez, S.E., Inlay, M.A., and Serwold, T. (2015). CD201 and CD27 identify hematopoietic stem and progenitor cells across multiple murine strains independently of Kit and Sca-1. Experimental Hematology 43, 578–585.

Visser, J.W., Bol, S.J., and van den Engh, G. (1981). Characterization and enrichment of murine hemopoietic stem cells by fluorescence activated cell sorting. Experimental Hematology 9, 644–655.

Walter, D., Lier, A., Geiselhart, A., Thalheimer, F.B., Huntscha, S., Sobotta, M.C., Moehrle, B., Brocks, D., Bayindir, I., Kaschutnig, P., et al. (2015). Exit from dormancy provokes DNA-damage-induced attrition in haematopoietic stem cells. Nature 520, 549–552.

Wilson, A., Laurenti, E., Oser, G., van der Wath, R.C., Blanco-Bose, W., Jaworski, M., Offner, S., Dunant, C.F., Eshkind, L., Bockamp, E., et al. (2008). Hematopoietic Stem Cells Reversibly Switch from Dormancy to Self-Renewal during Homeostasis and Repair. Cell 135, 1118–1129.

Wu, X., Dao Thi, V.L., Huang, Y., Billerbeck, E., Saha, D., Hoffmann, H.-H., Wang, Y., Silva, L.A.V., Sarbanes, S., Sun, T., et al. (2018). Intrinsic Immunity Shapes Viral Resistance of Stem Cells. Cell 172, 423–438.e425.

Yamamoto, R., Morita, Y., Ooehara, J., Hamanaka, S., Onodera, M., Rudolph, Karl L., Ema, H., and Nakauchi, H. (2013). Clonal Analysis Unveils Self-Renewing Lineage-Restricted Progenitors Generated Directly from Hematopoietic Stem Cells. Cell 154, 1112–1126.

